# Acute Myeloid Leukemia Driven by the CALM-AF10 Fusion Gene is Dependent on BMI1

**DOI:** 10.1101/524066

**Authors:** Karina Barbosa, Anwesha Ghosh, Anagha Deshpande, Bo-Rui Chen, Younguk Sun, Marla Weetall, Scott A. Armstrong, Stefan K. Bohlander, Aniruddha J. Deshpande

## Abstract

A subset of acute myeloid and lymphoid leukemia cases harbor a t(10;11)(p13;q14) translocation resulting in the CALM-AF10 fusion gene. Standard chemotherapeutic strategies are often ineffective in treating patients with CALM-AF10 fusions. Hence, there is an urgent need to identify molecular pathways dysregulated in CALM-AF10-positive leukemias which may lay the foundation for novel targeted therapies. Here we demonstrate that the Polycomb Repressive Complex 1 gene *BMI1* is consistently overexpressed in adult and pediatric CALM-AF10-positive leukemias. We demonstrate that genetic *Bmi1* depletion abrogates CALM-AF10-mediated transformation of murine hematopoietic stem and progenitor cells (HSPCs). Furthermore, CALM-AF10-positive murine and human AML cells are profoundly sensitive to the small-molecule BMI1 inhibitor PTC209 as well as to PTC596, a compound in clinical development that has been shown to result in downstream degradation of BMI1 protein. PTC-596 significantly prolongs survival of mice injected with a human CALM-AF10 cell line in a xenograft assay. In summary, these results validate BMI1 as a *bonafide* candidate for therapeutic targeting in AML with CALM-AF10 rearrangements.

## INTRODUCTION

Acute leukemia patients often harbor genomic translocation events that give rise to oncogenic fusion proteins (Greaves & Wiemels, 2003; Rowley, 1999). The t(10;11)(p13;q14) translocation is a recurrent, balanced translocation observed in human leukemia, which gives rise to the CALM-AF10 fusion protein (Bohlander et al., 2000; D Caudell & Aplan, 2008). Patients harboring the CALM-AF10 fusion have a particularly poor prognosis (Dreyling et al., 1998; Narita et al., 1999). Standard chemotherapeutic strategies are often not very effective in treating patients with CALM-AF10 fusions. Hence, there is an urgent need to identify molecular pathways dysregulated in CALM-AF10 positive leukemias which may lay the foundation for novel targeted therapies.

The N’-terminal partner of the fusion - CALM/PICALM, is a ubiquitously expressed component of the clathrin-mediated endocytosis (CME) pathway (Tebar, Bohlander, & Sorkin, 1999). Mutations in the murine *Picalm* gene are associated with defects in iron uptake and hematopoiesis (Klebig et al., 2008). CALM deletion in the hematopoietic system leads to severe deficiencies in endocytic vesicle formation, transferrin-mediated iron uptake and erythropoiesis (Ishikawa et al., 2015; Scotland et al., 2012; Suzuki et al., 2012). The C’-terminal fusion partner AF10 (MLLT10) on the other hand, is a PHD finger-containing chromatin reader protein that acts as a co-factor for the histone methyltransferase DOT1L (Chen et al., 2015; Aniruddha J. Deshpande et al., 2014). AF10 binds to the N-terminal histone H3 tail, with a preference for unmethylated lysine 27 (H3K27) (Chen et al., 2015). Methylation of H3K27 strikingly lowers the affinity of the N-terminal chromatin-reading PHD-zinc knuckle-PHD module (PZP) of AF10 for chromatin (Chen et al., 2015). Therefore, AF10 preferentially localizes to active chromatin domains where there are no repressive H3K27 methylation marks.

Early clues regarding the oncogenic mechanisms of CALM-AF10 came from transcriptome profiling studies using microarrays in AML and T-ALL. These studies showed that CALM-AF10-rearranged leukemias display a distinct gene expression signature (Dik et al., 2005a; Medhanie Assmelash Mulaw et al., 2008). This signature resembles the transcriptome of leukemia cells with rearrangements of the mixed lineage leukemia (MLL)-gene in terms of elevated expression of the posterior *HOXA* genes and the TALE-domain co-factor *MEIS1*. One striking difference between CALM-AF10-rearranged and MLL-rearranged AMLs was the consistently elevated expression of *BMI1* in CALM-AF10-rearranged cases. Elevated *BMI1* expression is observed in AML as well as T-acute lymphoblastic leukemia (T-ALL) with CALM-AF10 rearrangements (Dik et al., 2005a; Medhanie Assmelash Mulaw et al., 2008).

BMI1 is a member of the Polycomb Repressive Complex 1 (PRC1) with critical roles in the repression of developmentally important genes, including genes involved in the self-renewal of somatic stem cells (reviewed in Park, Morrison, & Clarke, 2004; Schuringa & Vellenga, 2010). The most well-documented role of *Bmi1* is in the epigenetic repression of the *Ink4a* locus genes *p16^Ink4a^* and *p19^Arf^*. This repressive activity of BMI1 is critical for its role in regulating cell-cycle progression and self-renewal of stem cells (reviewed in Park et al., 2004; Schuringa & Vellenga, 2010). BMI1 mediates this repressive activity on chromatin by stimulating the enzymatic activity of the PRC1 RNF2/RING2 E3 ligase (Cao, Tsukada, & Zhang, 2005). RNF2/RING2, the enzymatic component of the PRC1 complex, is responsible for the mono-ubiquitination of histone 2A at lysine 119 (H2AK119ub) leading to the epigenetic silencing of transcripts from H2AK119 monoubiquitylated promoters (Buchwald et al., 2006; Cao et al., 2005; Kallin et al., 2009).

High BMI1 expression is linked to oncogenic self-renewal in several tumors (reviewed in Park, Morrison, & Clarke, 2004; Schuringa & Vellenga, 2010). BMI1 overexpression is also implicated in epithelial to mesenchymal transition (EMT), metastasis and chemotherapy resistance in solid tumors (Siddique & Saleem, 2012) marking this gene as an attractive therapeutic target in several human malignancies.

In this study, we investigated the role of BMI1 in CALM-AF10-rearranged AML using genetic and pharmacological approaches. Our results, using mouse and human models, demonstrate that genetic or pharmacological BMI1 inhibition impairs CALM-AF10-mediated leukemogenesis *in vitro* and *in vivo*.

## RESULTS

### Genetic *Bmi1* deletion impedes CALM-AF10-driven myeloid transformation

We analyzed *BMI1* expression in RNAseq data from leukemia patients in the cancer genome atlas (TCGA, http://cancergenome.nih.gov/) as well as the recently reported pediatric pan-cancer genome alteration studies (Ma et al., 2018). We observed that patient samples with AF10-fusions, including both CALM-AF10 as well as MLL-AF10 fusions expressed significantly higher levels of *BMI1* compared to non AF10-rearranged samples. This was true for AML patients from TCGA studies as well as for childhood leukemia patients (AML, B-ALL, T-ALL and Mixed-lineage leukemia) from the pediatric pan-cancer studies (Ma et al., 2018) (Fig. S1A). These observations indicate that *BMI1* may be directly activated by AF10-fusion oncogenes as suggested previously (M. A. Mulaw et al., 2012).

Given the high-level expression of *BMI1* in AF10-rearrangements, we wanted to investigate the potential requirement for BMI1 in CALM-AF10-driven AML. Towards this end, we utilized a well-established model of retroviral CALM-AF10 overexpression in murine hematopoietic stem and progenitor cells (HSPCs) (A J Deshpande et al., 2011a). First, we tested whether *Bmi1* deficiency can affect CALM-AF10-mediated oncogenic HSPC transformation. We obtained mice in which the *Bmi1* locus is replaced by the GFP transgene (termed BKa.Cg-Ptprc^b^ *Bmi1^tm1Ilw^ Thy1*^a^/J mice). Since global *Bmi1* depletion leads to post-natal lethality in mice (Park et al., 2003), we isolated fetal liver cells from day 14.5 mouse embryos that were heterozygous or homozygous for the *Bmi1 null* allele. We transduced these cells or their *Bmi1* wild-type counterparts with a retroviral expression vector encoding a highly oncogenic version of the CALM-AF10 fusion oncogene (A. J. Deshpande et al., 2011) along with a bicistronic TdTomato fluorescent reporter. TdTomato+ cells were sorted by fluorescence-assisted cell sorting (FACS) and plated for colony-forming unit (CFU) assays according to the scheme in (Fig. 1A). We observed that CALM-AF10 transduced cells showed a significant decrease in their ability to form undifferentiated, blast-like colonies, upon loss of *Bmi1* alleles, while differentiated colony formation was not significantly impaired (Fig. 1B). These results indicate that Bmi1 is required for the immortalization of murine hematopoietic cells by the CALM-AF10 fusion gene. We then sought to determine whether hematopoietic cells already transformed by CALM-AF10 are also sensitive to *Bmi1* deletion. For this, we made use of another well-defined mouse with floxed *Bmi1* alleles (*Bmi1*^tm1.1Sjm^/J) which would allow for conditional ablation of the *Bmi1* gene. We immortalized HSPCs from these *Bmi^fl/fl^* mice with the CALM-AF10 fusion gene. Subsequently, we transduced these rapidly growing cells with a retrovirally encoded estrogen-receptor-fused Cre recombinase (ER-Cre) plasmid or a constitutive Cre-recombinase. Treatment with 4-hydroxytamoxifen (4-OHT) induced Cre-recombinase activity from the ER-Cre transduced cells, leading to excision of floxed *Bmi1* alleles (Fig. 1C and data not shown). We then performed *in vitro* proliferation as well as CFU assays from CALM-AF10 transformed bone marrow cells treated with 4-OHT or vehicle control (DMSO). We observed that *Bmi1* deletion led to a significant and progressive decline in viable cell numbers compared to vehicle-treated cells (Fig. 1D). This decrease in proliferation was accompanied by a significant increase in apoptotic cells as measured by Annexin V staining, as well as an increased ratio of cells in the G0/G1 compared to the S-phase (Fig. 1E). Using CFU assays, we also observed a significant decrease in the clonogenic capability of CALM-AF10-transformed cells upon *Bmi1* excision. Even though *Bmi1* deletion reduced the overall number of CFUs, the most striking reduction was observed in colonies with an undifferentiated or blast-like morphology (Fig.1F). Taken together, these experiments, using *Bmi1* constitutive or conditional knockout-mice, revealed that Bmi1 is critical for the initiation as well as maintenance of transformation by the CALM-AF10 fusion oncogene.

**Figure 1.**
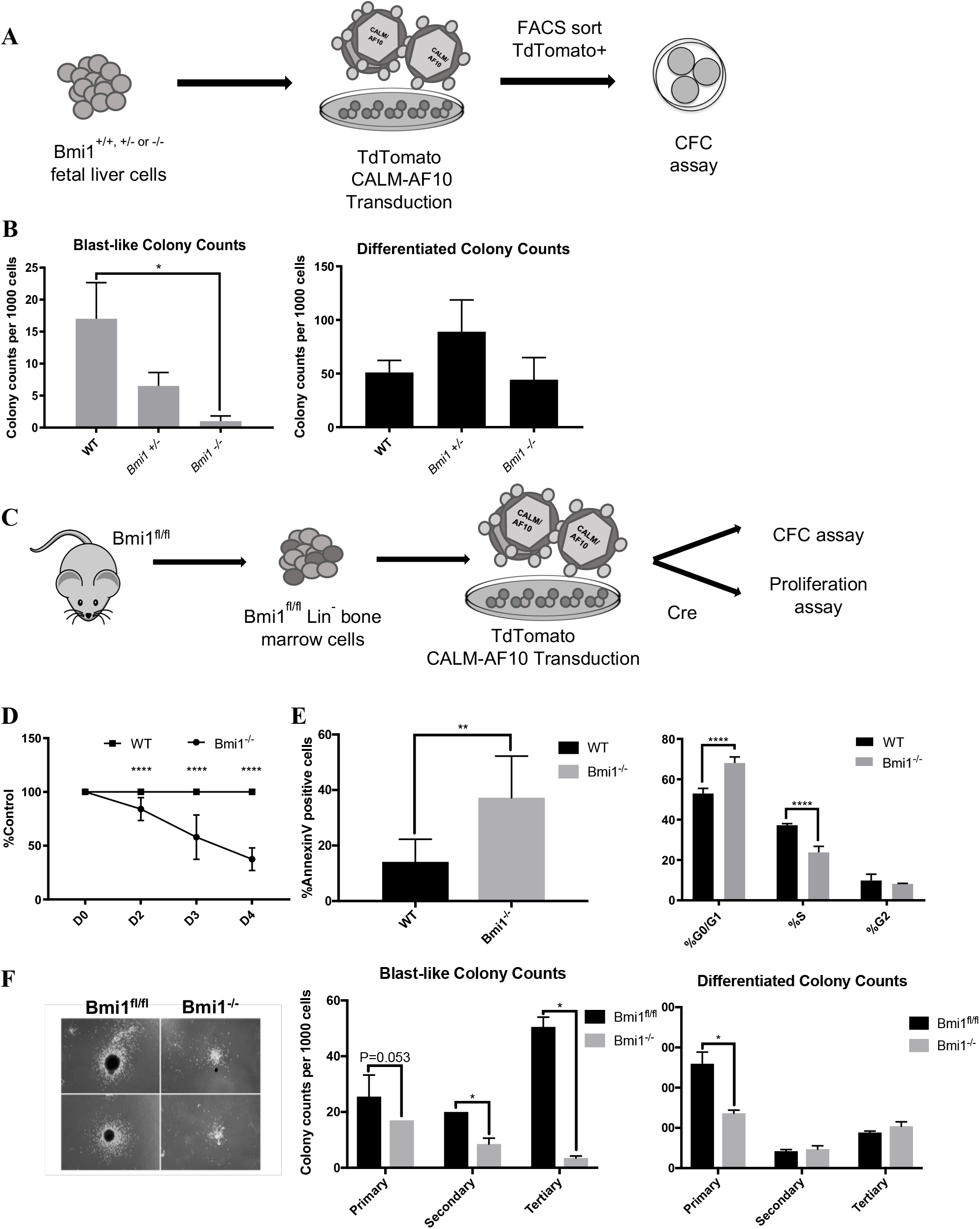
(A) Diagram illustrating the generation of retroviral CALM-AF10 transformed mouse cells with *Bmi1* wild-type or deficient backgrounds (fl/fl, fl/-, -/-). (B) Colony forming units of CALM-AF10 transduced mouse fetal liver *Bmi1* mutant cells with blast-like (left) or differentiated colonies (right) at day 7. *P<0.05, n=2 (C) Diagram illustrating the generation of CALM-AF10 transformed mouse cells with *Bmi1*^fl/fl^ backgrounds for *Bmi1* Cre-excision. (D) Analysis of *in vitro* cell proliferation of *Bmi1* excised mouse cells. *P<0.05, n=3. (E) (left) AnnexinV staining in *Bmi1* excised mouse cells. *P<0.05, n=3. (right) Cell-cycle progression analysis by propidium iodide staining in *Bmi1* deleted mouse cells. *P<0.05, n=3. (F) Representative images of colonies from CALM-AF10-transduced *Bmi1* deleted and wild-type cells 1 week after plating. (I) colony-forming units of CALM-AF10 transduced blast-like (left) or differentiated cells (right) at day 7. *P<0.05, n=2.

### Small-molecule BMI1 inhibition impairs murine CALM-AF10 AML growth and survival

Recently, small molecule inhibitors of BMI1 have been developed (Kreso et al., 2014; Nishida et al., 2017). We wanted to investigate whether these BMI1 inhibitors are active against CALM-AF10-driven AML. In order to do so, we utilized PTC-209, a novel small molecule inhibitor of BMI1, previously used to target colorectal cancer, chronic leukemia and multiple myeloma (Mayr, Neureiter, Wagner, Pichler, & Kiesslich, 2015; Mourgues et al., 2015). PTC-209 has been reported to reduce expression of BMI1 protein by altering regulation of the translation of the BMI1 mRNA (Kreso et al., 2014). We examined the effect of PTC-209 on mouse CALM-AF10 AML cell-growth by treating primary AML cells from three independently derived tumors. Cells were exposed to varying concentrations of PTC-209 for up to six days together with DMSO treated controls, and viable-cell counting performed every two days. First, we confirmed on-target activity of PTC-209 by ensuring the derepression of the Cdkn1a locus, a well characterized tumor suppressor locus that is transcriptionally repressed by Bmi1 activity (Fig. S2A). We then assessed the effect of PTC209 on CALM-AF10-transformed AML cells in proliferation assays *in vitro*. We observed that PTC209 induced a highly significant and concentration-dependent decrease in the number of viable CALM-AF10 AML cells over time (Fig. 2A) in comparison to their vehicle-treated counterparts. This decrease in cell viability was accompanied by a significant increase in AnnexinV positive cells, demonstrating induction of apoptosis upon PTC-209 treatment (Fig. 2B). The sensitivity of CALM-AF10 AML cells to PTC-209 was in the nanomolar range with an EC_50_ of 500nM.

**Figure 2.**
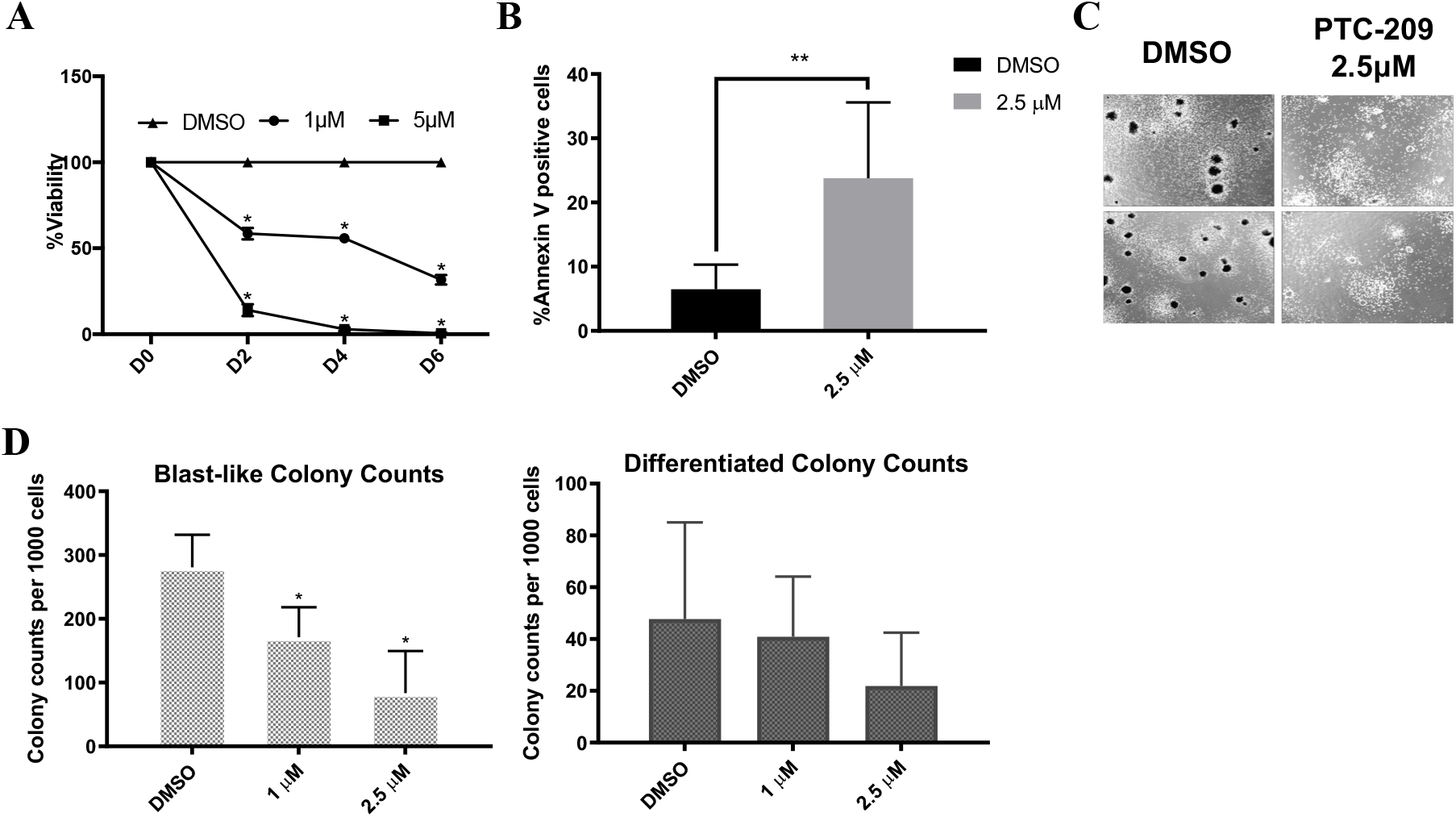
(A) Percentage of viable cells from mouse CALM-AF10-transformed AML cells upon PTC209 treatment compared to DMSO treated counterparts. *P<0.05, n=9. (B) Annexin V staining in mouse CALM-AF10-transformed cell lines. *P<0.05, n=9. (C) Representative images of colonies from mouse CALM-AF10-transformed cells upon 7 days of PTC-209 treatment. (D) Colony Formation Units of CALM-AF10 immortalized mouse cells upon PTC-209 vs DMSO treatment. Blast (left) or differentiated colonies (right) at day 7. *P<0.05, n=2.

Furthermore, compared to vehicle-treated AML cells, PTC-209 treatment also significantly reduced the clonogenic capacity of CALM-AF10 cells in CFU-assays in a concentration-dependent manner (Fig. 2C and 2D). These results demonstrate that small-molecule BMI1 inhibitor significantly inhibits the proliferative activity as well as clonogenic capability of murine CALM-AF10 AML cells.

### Pharmacological BMI1 inhibition impairs human CALM-AF10 AML *in vitro* and *in vivo*

We wanted to confirm our findings from the mouse models in human CALM-AF10-rearranged AML. For this, we treated the CALM-AF10-rearranged AML cell lines U937, KPMOTS and P31/Fujioka *in vitro* with DMSO or PTC209 at various concentrations, and assessed the effect on proliferation, cell cycle and apoptosis. In proliferation assays, we observed a significant and concentration-dependent decrease in viable cell counts of all three cell lines upon treatment with PTC209, compared to DMSO controls (Fig. 3A). PTC-209 also induced consistent increases in Annexin V positive cell populations compared to the vehicle, with the most significant and most pronounced difference in P31/Fujioka cells (Fig. 3B). Similar to mouse CALM-AF10-transformed cells, BrdU incorporation analysis showed an increase in the proportion of G0/G1 phase cells in U937, P31/Fujioka and KPMOTS cell lines, with a corresponding decrease in S-phase cells (Fig. 3C). Taken together, small-molecule BMI1 inhibition results in cell-cycle arrest features that are coupled with increased cell death and reduced proliferation of CALM-AF10-driven human AML cell lines, in concordance WITH our mouse model data.

**Figure 3.**
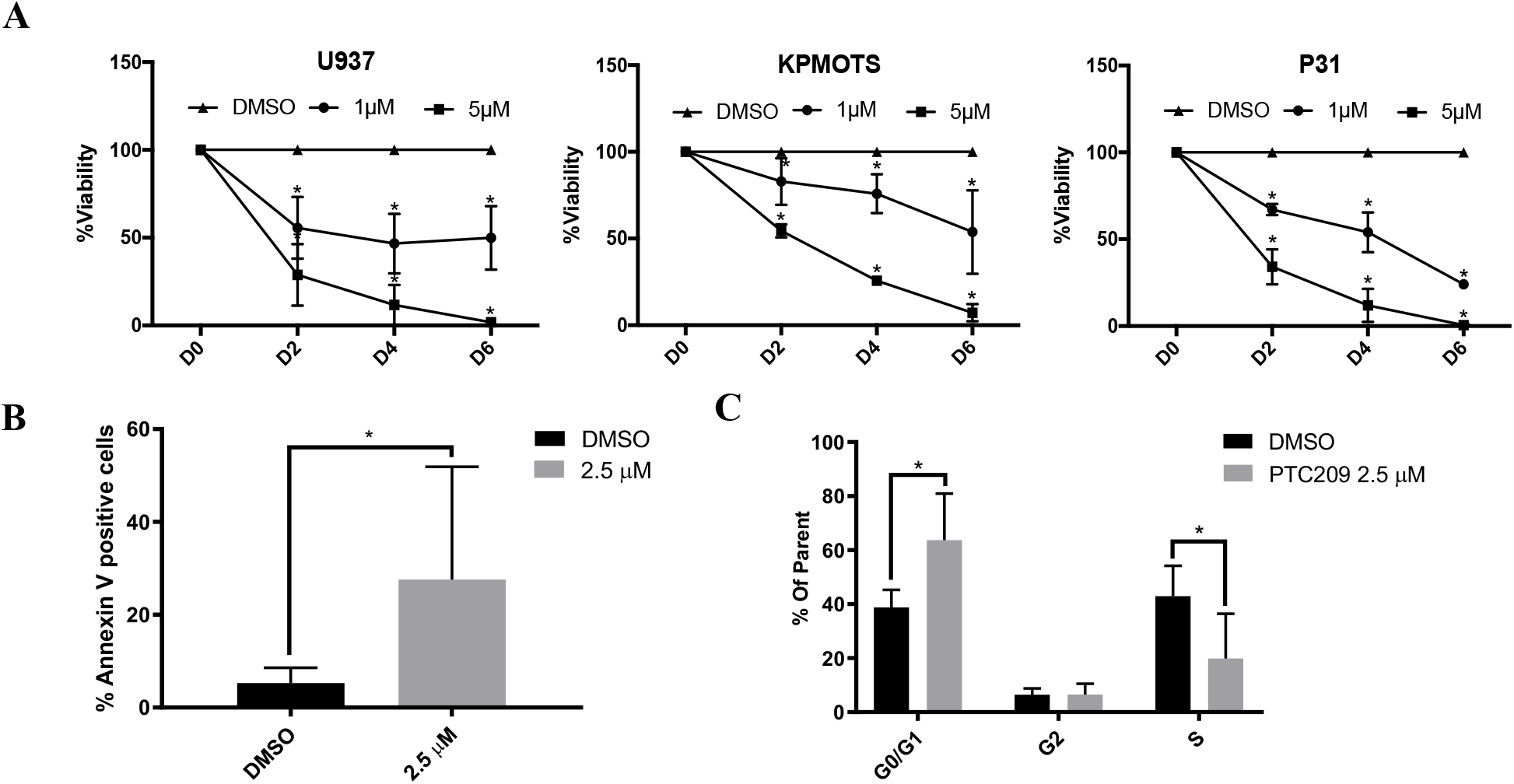
(A) Analysis of *in vitro* cell proliferation of human CALM-AF10-driven cell lines upon PTC209 treatment. *P<0.05, n=6. (B) AnnexinV staining in human CALM-AF10-driven cell lines. *P<0.05, n=3. (C) Cell-cycle analysis by BrdU incorporation in human CALM-AF10-driven cell lines is shown. *P<0.05, n=3.

Next, we wanted to validate our findings *in vivo*. For this, we used PTC596, a compound in clinical development identified by its ability to inhibit proliferation of BMI1 positive cancer stem cells (reference). The mechanism of BMI1 protein reduction is thought to be due to G2/M arrest causing accelerated ubiquitination and degradation of the BMI1 protein (reference). We assessed the ability of PTC596 to inhibit CALM-AF10-driven *in vivo* leukemogenesis. First, we injected immunodeficient mice (NRG-SGM3) with the P31/Fujioka cell line. Ten days after injection, we confirmed engraftment of P31 cells in mice by flow-cytometric assessment of the human CD45 marker (data not shown). We then orally administered one cohort of mice with PTC-596 and an age and engraftment matched cohort was administered the vehicle control. We observed that PTC596-treatment significantly delayed the latency of disease in mice compared to controls (Fig. 4) demonstrating the *in vivo* efficacy of small-molecule BMI targeting in this setting.

**Figure 4.**
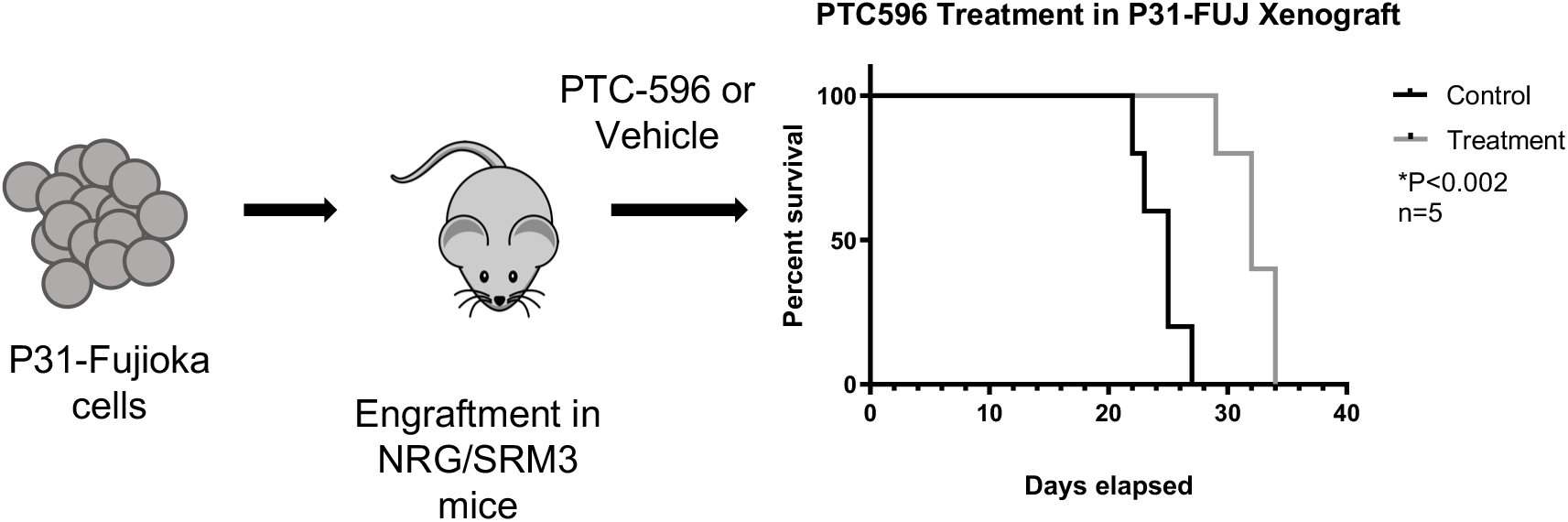
Diagram illustrating *in vivo* human CALM-AF10 cell line engraftment model and treatment with clinical-grade inhibitor PTC596. Survival curve for PTC596 vs. vehicle treated animals is shown on the right (n=5 mice per group, *P<0.002).

In summary, our results demonstrate that BMI1 is a *bonafide* candidate for therapeutic targeting in AML with CALM-AF10 rearrangements and possibly other leukemias with CALM-AF10 rearrangements.

## MATERIALS AND METHODS

### Reagents

PTC-209 was obtained from Cayman Chemical Co. (Ann Arbor, MI, USA), dissolved in DMSO and stored at −80 °C. PTC596 and its vehicle solution for *in* vivo studies were provided by PTC Therapeutics (South Plainfield, NJ, USA).

### Animal experiments

*Bmi1*-green fluorescent protein (GFP) transgenic mice (BKa.Cg-*Ptprc^b^ Bmi1^tm1Ilw^ Thy1*^a^/J) mice were obtained from The Jackson Laboratories (Bar Harbor, ME, USA, JAX # 017351) and maintained in the SBP animal facility. *Bmi1^fl/fl^* (Mich et al., 2014) mice were obtained from the Jackson Laboratories (JAX #028974). All experiments using mice were conducted as per procedures approved by the SBP Institutional Animal Ethics Committee.

### Cell culture

The human acute myeloid leukemia (AML) cell line U937 was a kind gift of Daniel Tenen, Beth Israel Deaconess Medical Center. The P31/Fujioka cells were obtained from the JCRB cell bank (#JCRB0091). KP-MO-TS was a kind gift from Dr. Issay Kitabayashi, National Cancer Center, Tokyo. All cell lines were cultured in RPMI-1640 (Gibco, Grand Island, NY, USA) medium supplemented with 10% heat-inactivated fetal bovine serum (FBS), 2 mM L-glutamine (Gibco) and 100 U/ml penicillin/streptomycin (Gibco).

Murine bone marrow cells from the femur and tibia were depleted of lineage positive cells using the EasySep™ Mouse Hematopoietic Progenitor Cell Isolation Kit (#19856, StemCell Techologies, Vancouver, Canada) and cultured overnight in DMEM (Gibco) media supplemented with 15% FBS (Gibco), 2 mM L-glutamine (Gibco), 100 U/ml penicillin/streptomycin (Gibco)and a cytokine cocktail containing mIL3, mIL6 and mSCF (Peprotech Inc). The following day, the cells were transduced with a retrovirally encoded version of CALM-AF10 bearing the C-terminal clathrin-binding domain of CALM and the octapeptide motif-leucine-zipper of AF10, as described in A J Deshpande et al., 2011. These BM progenitors transduced with the CALM-AF10 fusion were then either used directly for experiments or injected into mice.

### *In vivo* drug experiment

The xenograft model was established with NRG/SRM3 mice obtained from the SBP Animal Facility. Mice were irradiated (2.5 Gy) and 2×10^5^ P31/Fujioka cells were then injected intravenously. At 9 days post-injection, mice were randomized into two groups (n=5) to receive either PTC596 (12.5 mg/kg) or vehicle (0.5% hydroxypropyl methylcellulose and 0.1% Tween 80 in distilled water) by oral gavage twice per week. Circulating leukemia cells were detected on day 8 by flow cytometry using a human-specific CD45 antibody (#404012, Biolegend, San Diego). Mice were sacrificed upon signs of morbidity.

### Embryo generation and isolation of fetal liver cells

Embryos were generated from timed matings between male and female BKa.Cg-Ptprcb Bmi1tm1Ilw Thy1a/J mice. Detection of the vaginal plug was designated as E0.5. Pregnant females were sacrificed by cervical dislocation at E14.5. The uterine horns were removed and fetuses were separated from maternal tissue. Fetal livers were dissected, and single cell suspensions were obtained by straining tissues through a 10-micron mesh. Red blood cell (RBC) lysis was performed using 1X RBC Lysis Buffer (Sigma-Aldrich) according to the manufacturer’s guidelines. Fetal liver cells were cultured in DMEM (Gibco, Grand Island, NY) media containing 15% FBS, 2 mM L-glutamine (Gibco, Grand Island, NY), 100 U/ml penicillin/streptomycin (Gibco, Grand Island, NY) and a cytokine cocktail (mIL6, mIL3 and mSCF from Peprotech Inc, Rocky Hill, USA, #216-16, #213-13, #250-03).

### Cell proliferation assays

Mouse and human (U937, P31-Fujioka and KP-MO-TS) CALM-AF10 cell lines were seeded and treated with DMSO (control) or PTC-209 (1 μM or 5 μM). Samples were taken on the second, fourth and sixth day after setup of the assay. Cell viability on treatment of PTC-209 was determined by using Sytox Blue Dead Cell Stain (Invitrogen) via flow cytometry. Analysis was performed on an LSR Fortessa (BD Biosciences).

### Cell cycle and apoptosis assay

Apoptosis induction was determined by combined Annexin V and Sytox staining. CALM-AF10 cell lines were treated with either DMSO or PTC-209 at 2.5 μM. After 48h, cells were harvested and stained with Annexin V-APC (#550474, BD Pharmingen) for 15 minutes on ice in the dark. The cells were washed and stained with and Sytox Blue Dead Cell Stain (Invitrogen, Thermo Fisher Scientific) before performing analysis. Cell cycle assessment was performed after 20 min of labeling with a BrdU-APC Flow kit (#552598, BD Pharmingen), following the manufacturer’s guidelines. All flow cytometric analyses were performed on an LSR Fortessa (BD Biosciences).

### Colony formation assay

Mouse CALM-AF10 cell lines either treated or untreated with PTC-209 at 1 μM or 2.5 μM were plated in duplicates in 1.1 ml methylcellulose-based medium (MethoCult 3234, StemCell Techologies, Vancouver, Canada) per well, containing 460 pM mIL6, 1090 pM mSCF and 662 pM mIL3 (Peprotech Inc) and incubated for 7 days. At the end of the incubation period, the number of blast and differentiated colonies was scored using an inverted microscope.

### Quantitative RT-PCR

Total RNA was isolated using TRIzol (#15596026, Thermo Fisher Scientific, San Jose, CA, USA) according to the manufacturer’s instructions. cDNA was synthesized from extracted RNA using the ProtoScript First Strand cDNA Synthesis Kit (#E6300, New England Biolabs Inc, Beverly, MA, USA). CDKN1A expression levels were measured using quantitative PCR (qPCR) using TaqMan Universal PCR Master Mix and pre-designed TaqMan gene expression assays (Applied Biosystems, Foster City, CA, USA) on the Stratagene MX3000P (Agilent Technologies, La Jolla, CA, USA).

### Statistical Analysis

Flow cytometry data was analyzed using FlowJo (FlowJo Software, Tree Star, Ashland, OR). All statistical analyses were performed using GraphPad Prism 7 Software (San Diego, CA, USA), except for violin plots and analysis from Figure S1 for which the R Statistical Software was used. P values were calculated using the Student’s t-test.

## DISCUSSION

Despite the well-described oncogenic activity of BMI1 in a wide range of human cancers, therapeutic BMI1 targeting has proven to be elusive. Recently, the BMI1 small-molecule inhibitor PTC209 was developed by PTC Therapeutics (Kreso et al., 2014) and has been used in preclinical studies to treat chronic and acute myeloid leukemia (Mourgues et al., 2015), biliary tract cancer (Mayr et al., 2015), multiple myeloma (Bolomsky et al., 2017; Bolomsky, Schlangen, Schreiner, Zojer, & Ludwig, 2016), non-small cell lung cancer (Yong et al., 2016), glioblastoma (Jin et al., 2017), ovarian cancer(Dey et al., 2016), prostate cancer (Bansal et al., 2016), breast cancer (Dimri et al., 2016) and colorectal cancer (Kreso et al., 2014). Furthermore, a next-generation clinical-grade BMI1 inhibitor PTC596 has been developed, which is currently in clinical trials for advanced solid tumors (Infante et al., 2017). The fact that elevated *BMI1* expression is a common feature of CALM-AF10 leukemias regardless of lineage suggests that BMI1 may be a transcriptional target of the CALM-AF10 fusion protein. Intriguingly, we have previously noted that BMI1 is located on chromosome 10 adjacent and downstream of the wild-type AF10 gene (M. A. Mulaw et al., 2012). Therefore, it is also likely that the CALM-AF10 fusion-event drives enhanced BMI1 expression through the disruption of topologically associated domains (TADs) and juxtaposition of BMI1 to the strong, CALM-associated enhancers, as a result of the t(10;11) translocation. Since elevated *Bmi1* expression can be observed even in mouse models of CALM-AF10-driven AML (M. A. Mulaw et al., 2012) which do not harbor a t(10;11) translocation, the former scenario is more likely, although the latter possibility with TAD activation cannot completely be ruled out. Regardless of the mechanism of BMI1 activation, this characteristic CALM-AF10-associated molecular event may create a novel dependency that is therapeutically tractable.

The role of BMI1 in AML has been studied in the context of leukemias driven by other leukemia-associated oncogenes. Interestingly, genetic experiments demonstrate that the dependency on BMI1 is selective. Bmi1 was first reported to be important for AML stem cells in a murine study using a retroviral model of Hoxa9 and Meis1 co-expression (Lessard & Sauvageau, 2003). This study showed that *Bmi1* deletion does not affect the initiation of Hoxa9-Meis1-driven AML, but significantly impairs the ability of primary leukemias to transmit disease in secondary recipients (Lessard & Sauvageau, 2003). This study indicated that Bmi1 was important for the self-renewal of leukemia stem cells in the retroviral Hoxa9-Meis1 coexpression model. Other studies demonstrated that while myeloid transformation driven by the fusion oncoproteins AML1-ETO and PLZF-RARA is strongly sensitive to Bmi1 depletion (Boukarabila et al., 2009; Smith et al., 2011), leukemias driven by the MLL-AF9 fusion oncoprotein are not dependent on *Bmi1* expression (Smith et al., 2011). Another study with the MLL-AF9 fusion protein observed that *Bmi1* was necessary for the generation of AML from granulocyte macrophage progenitors (GMPs), indicating that BMI1 may be critical for leukemic transformation of downstream hematopoietic progenitors by MLL-AF9, but not for the transformation of hematopoietic stem cells (HSCs) (Yuan et al., 2011). These studies demonstrate the selective requirement of BMI1 depending on the mutational sub-type of AML as well as on the developmental stage of the leukemia cells. In our studies, CALM-AF10-driven transformation seems to require Bmi1 both for the initiation of transformation as well as maintenance. More recently, the small-molecule BMI1 inhibitors PTC209 as well as PTC596 have been used to demonstrate that a broad panel of human AML cell lines are sensitive to small-molecule BMI1 inhibition (Mourgues et al., 2015; Nishida et al., 2015, 2017) although none of these studies focused specifically on CALM-AF10-rearranged AML. In CALM-AF10, the clinical rationale for BMI1 targeting may be clearer, given that BMI1 is strongly upregulated in CALM-AF10 rearranged AML.

Our observation that CALM-AF10 fusions require BMI1 for initiation as well as maintenance of transformation provides pre-clinical evidence that pharmacological BMI1 inhibition may provide potential clinical benefit in leukemias with CALM-AF10 rearrangements. These results could help inform future clinical trials with BMI1 inhibitors. It is pertinent to note that BMI1 upregulation is observed not only in CALM-AF10 positive AML, but also in T-ALL, where elevated BMI1 expression was first noted (Dik et al., 2005b). It is therefore very likely that small-molecule BMI1 inhibition is a lineage-independent vulnerability associated with CALM-AF10-rearrangements. The hypothesis that CALM-AF10-rearranged T-ALLs may also be sensitive to BMI1 inhibition needs to be validated using T-ALL models of this disease. Even though CALM-AF10 fusions are more frequent in T-ALL than in AML, CALM-AF10 mouse models reported so far are biased towards the myeloid lineage (David Caudell, Zhang, Yang, & Aplan, 2007; A. J. Deshpande et al., 2011; Aniruddha J. Deshpande et al., 2006; Dutta et al., 2016), hampering the validation of therapeutic candidates for CALM-AF10 positive T-ALL. Future studies will focus on the development of CALM-AF10 T-ALL models and the validation of BMI1 inhibitors in these studies.

## AUTHOR CONTRIBUTIONS

KB, AD, BC, YS, and AG performed experiments, analyzed data and participated in manuscript preparation. AJD, SAA, MW and SKB participated in data analysis, expert advice or overall experimental design. SKB and AJD conceptualized the project, designed experiments and assisted in manuscript preparation.

## ACKNOWLEDGMENTS

We would like to acknowledge Yoav Altman and Amy Cortez at the SBP Flow cytometry core facility for sorting our samples and Buddy Charbono at the Animal Facility at SBP for mouse injections. We would like to acknowledge the support of the Lady Tata Memorial Foundation to A.D. We would like to acknowledge our funding sources: NIH R00 CA154880, NIH/NCI P30 CA030199, as well as the ASH Scholar award and the V-Foundation Award to A.J.D. S.K.B. was supported by Leukaemia and Blood Cancer New Zealand and the family of Marijana Kumerich.

## CONFLICTS OF INTEREST

AJD is a consultant at A2A Pharmaceuticals, New Jersey and Salgomed Therapeutics, La Jolla. MW is employed by PTC Therapeutics and has received salary and compensation for time, effort, and hold or held financial interest in the company.

---

**Supplementary Figure 1.**
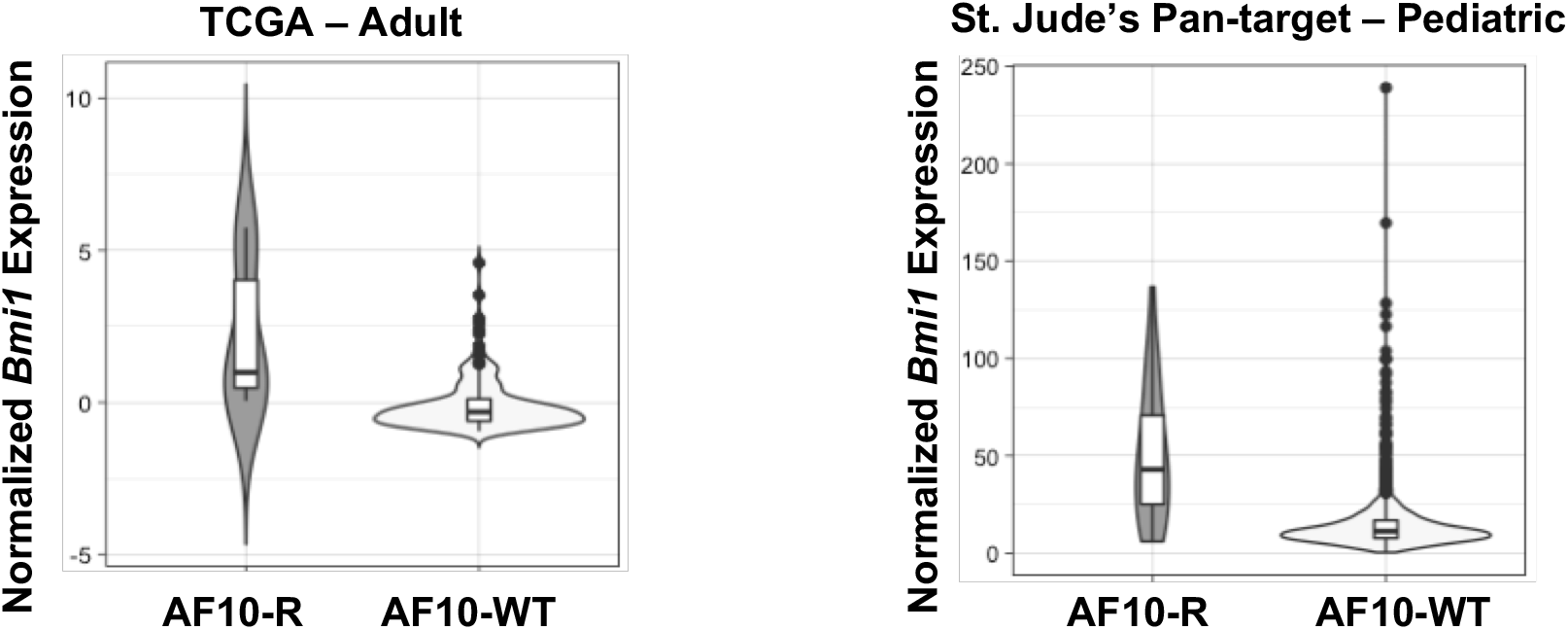
*BMI1* expression is plotted based on RNA-seq datasets from The Cancer Genome Atlas (TCGA LAML, left) or the Pediatric Cancer Data Portal (PeCan hematological malignancies, right). Patients harboring AF10 rearrangements (AF10-R, n=6 and n=27) are compared to patients without AF10 rearrangements (n=167 and n=1202). P value for PeCan comparison: 2e-16, for TCGA: 1.68e-5.

**Fig. S2.**
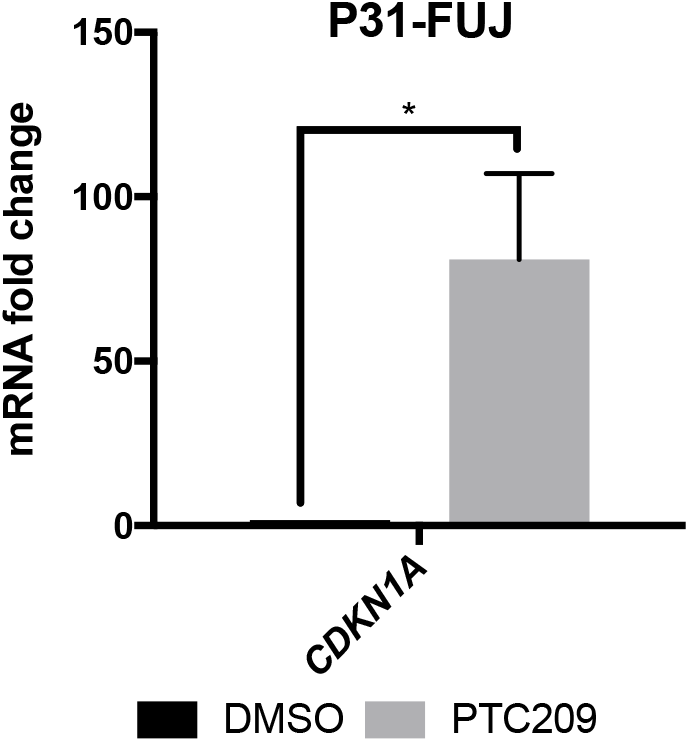

